# Synchronous and asynchronous counterillumination by three types of photophores in the firefly squid, *Watasenia scintillans*

**DOI:** 10.1101/2025.10.29.685447

**Authors:** Satoshi Kusama, Eiji Fujiwara, Osamu Inamura, Kenji Ishii, Miwa Tamura-Nakano, Noritaka Hirohashi

**Author notes:** These authors contributed equally (Satoshi Kusama, Eiji Fujiwara). To whom correspondence may be addressed. Corresponding author: Noritaka Hirohashi. **Author Contributions:** S.K., E.F., and N.H. designed research; S.K., E.F., M.T.N., and N.H. performed research; O.I. and K.I. contributed reagents/analytic tools; M.T.N. and N.H. analyzed data; and N.H. wrote the paper. **Competing Interest Statement:** The authors declare no competing interests.

## Abstract

Countershading—animal body coloration that is darker on the dorsum than the ventrum—is a form of crypsis achieved through a gradient in skin pigmentation. Counterillumination is somewhat similar, achieving crypsis in midwater marine organisms by emitting downward-directed bioluminescence. Although mesopelagic squids are assumed to use counterillumination, high-resolution spatiotemporal changes in illumination, particularly for each photophore type, have not been elucidated. *Watasenia scintillans* has three types of photophores on its ventral surfaces: blue- and green-emitting ventral photophores and ocular photophores. It is demonstrated here that all three types of photophores are synchronously, reversibly, and repeatedly involved in counterillumination in response to changes in dim overhead lighting. Furthermore, the green-emitting photophores maintained luminescence in darkness, and their on/off switching is precisely regulated within the ambient temperature range. The photoreceptors responsible for counterillumination are extraocular photosensitive vesicles (EOPs) rather than eyes. Exposure of EOPs to visible light (blue, green, and yellow) elicited counterillumination in proportion to light intensity. No color shifts (such as from green to blue) were observed in any of the photophores. While synchronous illumination seems to support the crypsis hypothesis, the ability to independently control each photophore type could also support a previous hypothesis of conspecific signaling.

**Significance Statement:** Bioluminescence is widespread among deep-sea organisms, yet the specific functions of individual light-producing organs distributed across the body surface remain underexplored. In this study, three types of ventral photophores in the firefly squid, *Watasenia scintillans*, were video-recorded, revealing their synchronous involvement in counterillumination—a mechanism that obscures the body silhouette when viewed from below against blue downwelling surface light. Notably, green-emitting photophores remained illuminated for extended periods in darkness, indicating a potential role in visual communication among conspecifics. The video-based techniques employed to quantify individual photophore activity provide new opportunities to investigate photo-signaling in animals inhabiting the mesopelagic zone.

## Introduction

Midwater pelagic nektonic and some planktonic animals face the inevitable risk of predation because there is nowhere to hide from visual predators. In particular, organisms are vulnerable to predators when targeted from below, as they appear as dark silhouettes against the background of downwelling light coming from the surface (1–4). To mitigate this effect, many mesopelagic organisms produce bioluminescence on their ventral surface at a brightness similar to that of their immediate surroundings (5, 6). The use of bioluminescence to break the silhouette when viewed from below is called counterillumination. However, to be effective and safe, the intensity of such ventrally directed bioluminescence needs to be finely tuned to match the downwelling light (7–13).

The Hawaiian midwater squids *Abraliopsis* sp. and *Abralia trigonura* are well known for their counterillumination, following experiments by Young and colleagues, who demonstrated a strong positive correlation between overhead brightness and bioluminescence intensity (14, 15). They also found that the color characteristics of the bioluminescence emitted by these squids shift from blue (deep-water-penetrating spectra) to green (moonlight spectra) as water temperature increases, corresponding to the conditions these species encounter during vertical migration in Hawaiian waters (16). The observed color change was suggested to occur through a spectral shift in at least one class of photophores, as well as through the recruitment of photophores with different wavelength-emission characteristics (16–18). Although it is thought that this color change is related to efficient camouflage (19–21), its mechanism remains largely elusive (but see (22, 23)). Furthermore, the findings of Young et al. are not without controversy in view of the following observations in the mesopelagic Japanese firefly squid, *Watasenia scintillans*: (1) Color changes in the photophores such as those reported by Young et al. have not been observed (24, 25); (2) there are distinct blue- and green-emitting photophores on the ventral surface (24–26); and (3) *W. scintillans* may use green bioluminescence for conspecific signaling (27, 28). This hypothesis was proposed following observations of the structural and spectrochemical characteristics of the eye, particularly the banked ventral retina, which receives light reaching the squid from above. This part of the retina is unusual because it consists of two pigmented layers (distal two-thirds yellow; proximal third pink), resulting in an estimated photosensitivity peak of ≈550 nm for the retinal cells in the deepest part of the ventral retina (27, 28). However, there have been no published reports regarding counterillumination in *W. scintillans* nor any other autogenic luminous squids aside from the Hawaiian squids reported by the studies of Young and colleagues. Questions for which answers are currently unknown are as follows. 1) Does counterillumination occur in *W. scintillans*? 2) If it does occur, does each type of photophore have its own distinct illumination specificity and sensitivity to environmental conditions such as brightness and temperature? 3) Do any of the photophores show any shifts in the wavelength of their light emission upon these environmental changes? Lastly, 4) Where are the photoreceptors that are responsible for counterillumination?

The present study used high-resolution colorimetric analysis of digital images obtained from an ultra-sensitive CMOS video camera to resolve the spatiotemporal dynamics of light emission under light-controlled conditions in captive firefly squids.

## Results

Under dim overhead lighting, *W. scintillans* individuals emit bioluminescence from the ventral surface (Fig. 1A). The ventral surface contains at least four distinct classes of photophores that differ in morphology, color and size. These include ocular photophores (oc-photo) located beneath each eye (Fig. 1B), as well as arrays of both blue and green photophores (G-photo) distributed across the entire ventral half (Figs. 1C-1G). Blue photophores can be categorized into two distinct sizes: large (B-photo) and small (b-photo). The colors of luminescence and of reflected light from the photophore surface illuminated by white LED light are similar (Figs. 1C, 1D). B-photo and b-photo are structurally indistinguishable (Figs. 1H, 1I). In contrast, oc-photo and G-photo exhibit distinct structural and histochemical characteristics (Figs. 1J, 1K, Figs. S1, S2). Bioluminescence was observed from all oc-photo (Fig. S3). However, they did not always emit light in synchrony, partly due to concealment by an overlay of dermal chromatophores and, otherwise, because they were apparently under independent control (Fig. S3; panels B, C).

**Figure 1.**
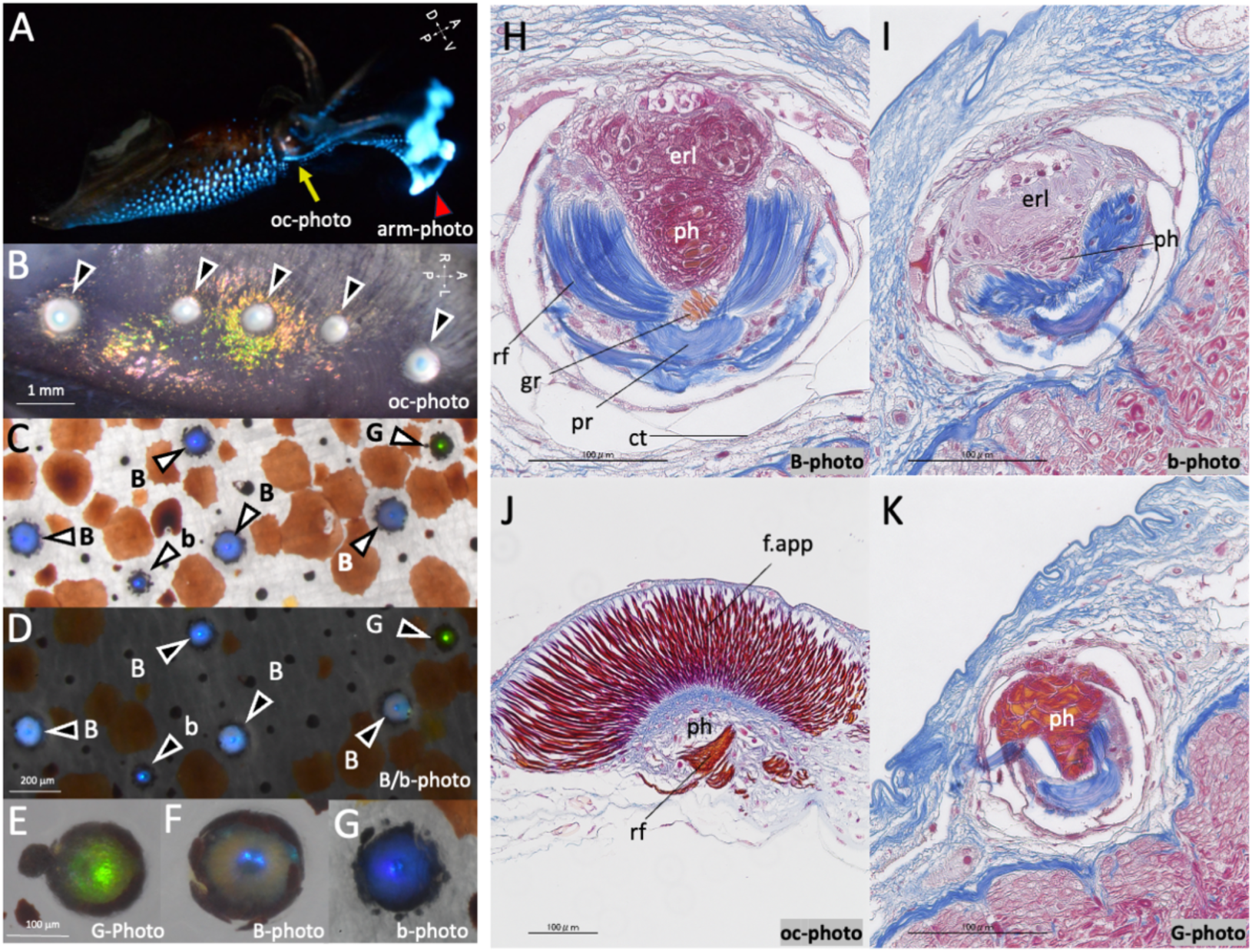
The ventrally located ocular photophores can emit light asynchronously in *Watasenia scintillans*. (A) Female *W. scintillans* displaying the counter-illuminating ventral body surface; the dorsal surface is dark with few photophores. The two most ventral pairs of arms on the right side are visible in right lateral view (orientation: A, anterior; D, dorsal; P, posterior; V, ventral). The saturated blue coloration at the right originates from large photophores at the tips of the most ventral arms (’arms IV’), which are not components of the counterillumination system but typically illuminate when the squid is startled or stressed in captivity. (B) Five ocular photophores (arrowheads) are attached to the ventral surface of the integument covering the right eyeball (orientation: A, anterior; L, left; P, posterior; R, right). (C, D) Bright-field (C) and dark-field (D) images of two additional photophores, one emitting green light (G) and the other blue (B/b). Blue photophores are distinguished by diameter as small (b) or large (B). (E-G) Bright-field images at higher magnification provide detailed views of the three main photophore types. (H-K) Cross-sections of B-photo (H), b-photo (I), oc-photo (J) and G-photo (K) stained with Masson-Trichrome. ph, photocytes; erl, endoplasmic reticulum lamellae (40); rf, reflector; gr, granules; pr, posterior reflector; ct, connective tissue; f-app, fibrous apparatus (26).

When maintained in captivity under aquatic conditions in darkness for 30 to 60 minutes, the animal remained unilluminated or displayed only minimal ventral illumination. Subsequently, all ventrally located photophores, including oc-photo, B/b-photo, and G-photo, exhibited counterillumination in response to exposure to a dim light (Figs. 2B, 2C, Movie S1-S3). The intensity and wavelength of bioluminescence emitted by individual photophores were estimated using pixel-wise color gamut calibration (Figs. 3A-3C; *Materials and Methods*). A simple linear approximation was applied to the relationship between light intensity and the RGB values on a logarithmic scale (Fig. 3B), as well as to the relationship between emission wavelength and ratiometric (B/G) color components using logarithmic axes (Fig. 3C). Using these standard calibrations, we observed a greater change in light intensity in B/b-photo (FC_on/off_: 6.09 ± 3.12, P < 0.001; FC_off/on_: 0.36 ± 0.20, P < 0.001; n = 18) and oc-photo (FC_on/off_: 4.20 ± 2.13, P < 0.001; FC_off/on:_ 0.475 ± 0.20, P < 0.001; n = 15),whereas emission from G-photo (FC_on/off:_ 2.12 ± 0.79, P < 0.001; FC_off/on:_ 0.93 ± 0.34, P = 0.24; n = 18) remained elevated longer in the dark (Figs. 3D, 3E). No significant shifts were observed in the estimated maximum wavelength of emission during the increase (Exact Wilcoxon signed rank test: G-photo, P = 0.724; B/b-photo, P = 0.068) or decrease (G-photo, P = 0.339; B/b-photo, P = 0.367) of bioluminescence in response to turning on an overhead light (Fig. 3F), suggesting that there is no substantial color shift between green and blue in the same photophores during counterillumination.

**Figure 2.**
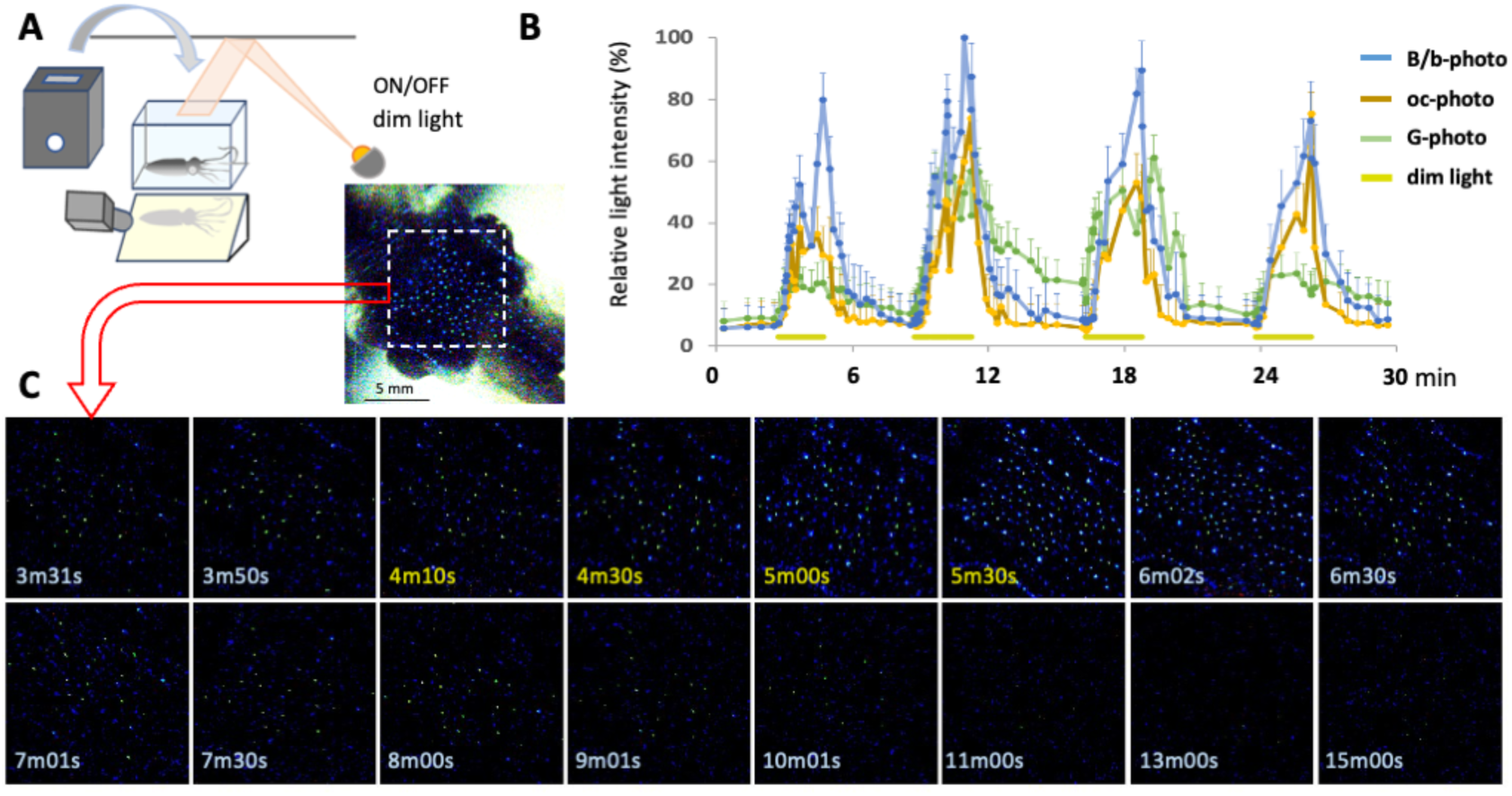
All three types of ventral photophores exhibit counterillumination in synchrony. (A) The experimental setup includes an overhead reflection lighting apparatus, a light-shielding container with a slot on top and a side hole for filming, an angled mirror that allows viewing of the bottom of the tank from a CMOS video camera mounted at the side of the tank. (B) Representative recording of illumination dynamics for each type of photophore showing repeated cycles of counterillumination in response to a dim light stimulus. (C) Representative series of video frames showing counterillumination photophore activity in response to overhead dim lighting (lights on during time = 4 to 6 min).

**Figure 3.**
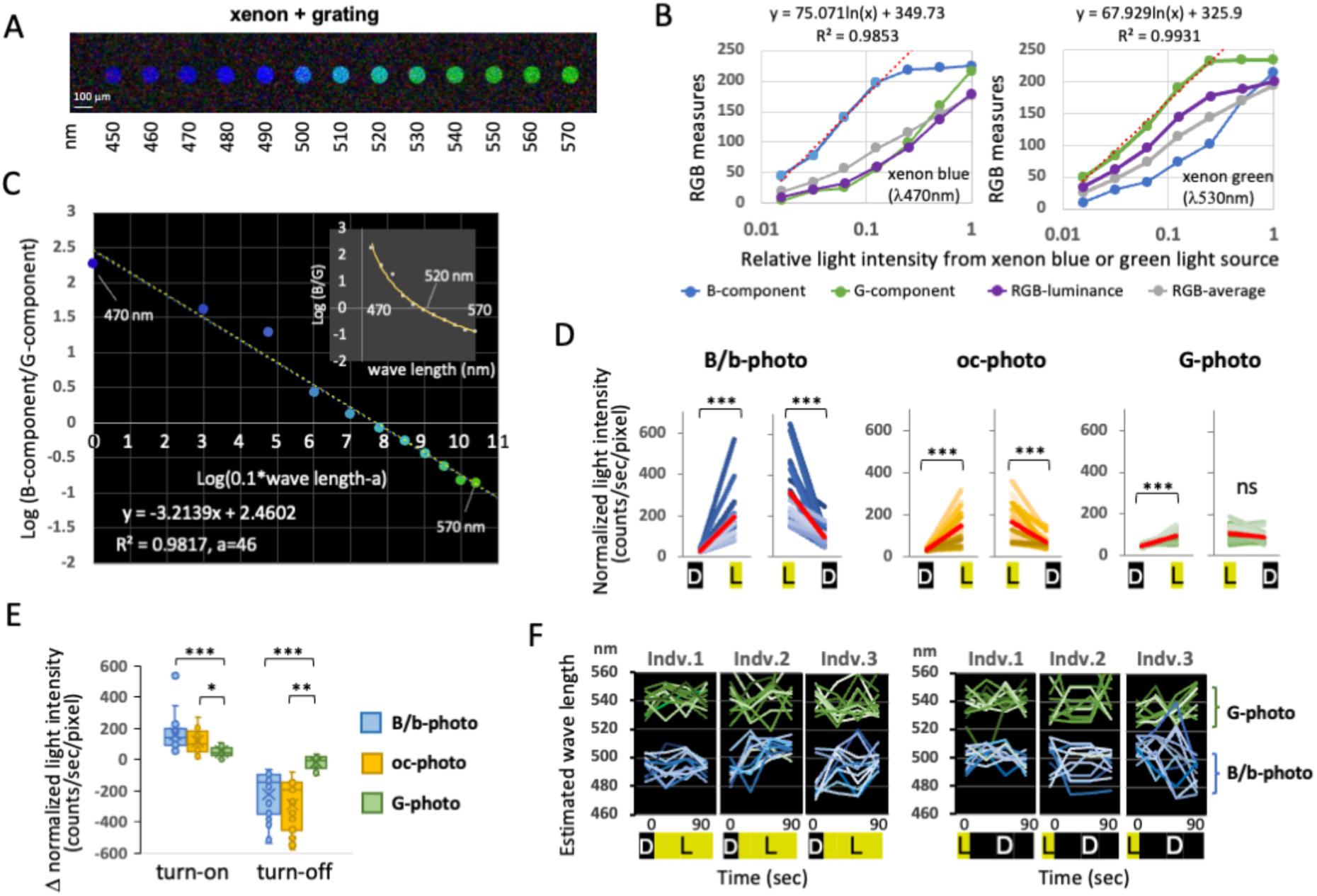
Calibration and estimation of light intensity and wavelength from video images of counterillumination. (A) Video images of standard peak spectrum (λmax) with narrow bandwidth generated by a diffraction grating of xenon light. (B) Standard calibration curves to estimate light intensity of xenon blue (λmax 470 nm; *left* panel) and xenon green (λmax 530 nm; *right* panel) after processing RGB measures of the still images extracted from video clips. (C) A log-log plot of ratiometric measurements (B-component/G-component) as a function of wavelengths, as shown in (A). (D, E) Reaction kinetics to the presence or absence of a dim light stimulus from overhead, based on measurements at 0 and 1 min after light on (D→L) or light off (L→D). Data from 15-18 photophores of four individuals. Mean values are indicated by red lines. F, Changes in estimated emission wavelengths of bioluminescence from each photophore (3 individuals) in 90-second durations after light on (D→L) or light off (L→D). Significance supported by Wilcoxon signed rank test (in D) or Mann–Whitney U-test (in E): *, p < 0.05; **, p < 0.01; ***, p < 0.001.

Next, the locations of photoreceptors responsible for counterillumination were investigated. A system was developed to enable spatiotemporal control of overhead illumination while simultaneously imaging both ventral and dorsal sides with supersensitive video cameras (Fig. 4A-4C). Counterillumination was observed immediately upon applying blue-light illumination to the entire dorsal side (Fig. 4D, none). In contrast, concealing only the head region eliminated counterillumination (Fig. 4D, head). However, when the head-shielding apparatus allowed light to reach the eyes and extraocular photosensitive vesicles (EOPs), counterillumination persisted (Fig. 4D, *headΔ^(eyes, EOPs)^*).

**Figure 4.**
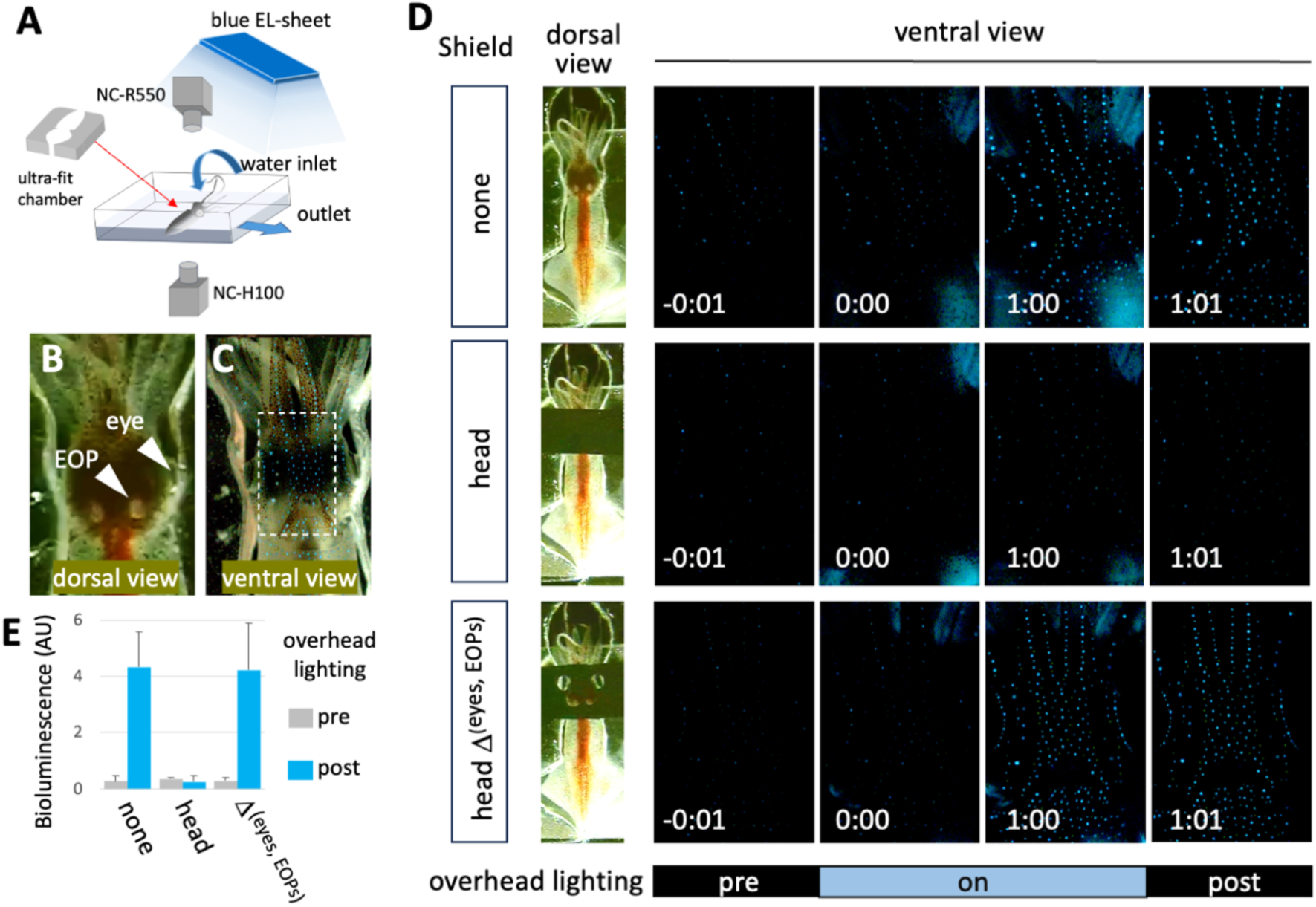
Identification of dorsally-localized photoreceptors for counterillumination. (A) A schematic diagram showing a ventral bioluminescence imaging system. The animal is immobilized in a semi-closed chamber (*ultra-fit chamber*) placed in a shallow-water container to restrict vigorous movements, while being freed for fin-beating and mantle contractions for breathing. The temperature-controlled water is circulated. Two cameras are mounted to capture simultaneous images of both the ventral and dorsal sides. (B, C) Still images of the head region from a video clip, showing dorsal (B) and ventral (C) views. *Arrowheads* point to the right eye (*eye*) and the right extraocular photosensitive vesicles (*EOP*). A *broken box* indicates the area of dark-field images shown in (D). (D) Time series of images showing the counterilluminating or non-illuminating ventral head region by turning the overhead blue light on during the time between 0:00 and 1:00 (*on*) from the dorsal side, with/without different types of head shields. The head shielding apparatus covers the entire head region (*head*) or allows light passage at points of eyes and EOPs (*headΔ^(eyes, EOPs)^).* Time code indicates min:sec. Data represent mean ± SEM (n = 4).

The system was further refined by integrating a pair of optical fibers guided by three-axis manipulators, enabling precise positioning of light exposure to the eyes or EOPs at selected wavelengths and intensities (Figs. 5A-5C, Fig. S4). When optical light at peak wavelengths of 488, 546, or 580 nm was directed to the right side of the EOPs, ventral illumination increased proportionally with light intensity (Figs. 5D, 5E, Fig. S4). In contrast, ventral illumination was rarely observed when 620 nm light was applied to the EOPs, when any wavelength was directed to an eye, or when 546 nm light was applied to both eyes (Figs. 5D-5H). The intensity of ventral illumination was sensitive to water temperature, reaching its maximum at approximately 9.0 °C (Fig. 5I, Fig. S5). The contribution of G-photo to overall ventral illumination was lowest under conditions of low light exposure or low temperature (5.0 °C) and increased as either light intensity or temperature rose (Figs. 5J, 5K, Fig. S6).

**Figure 5.**
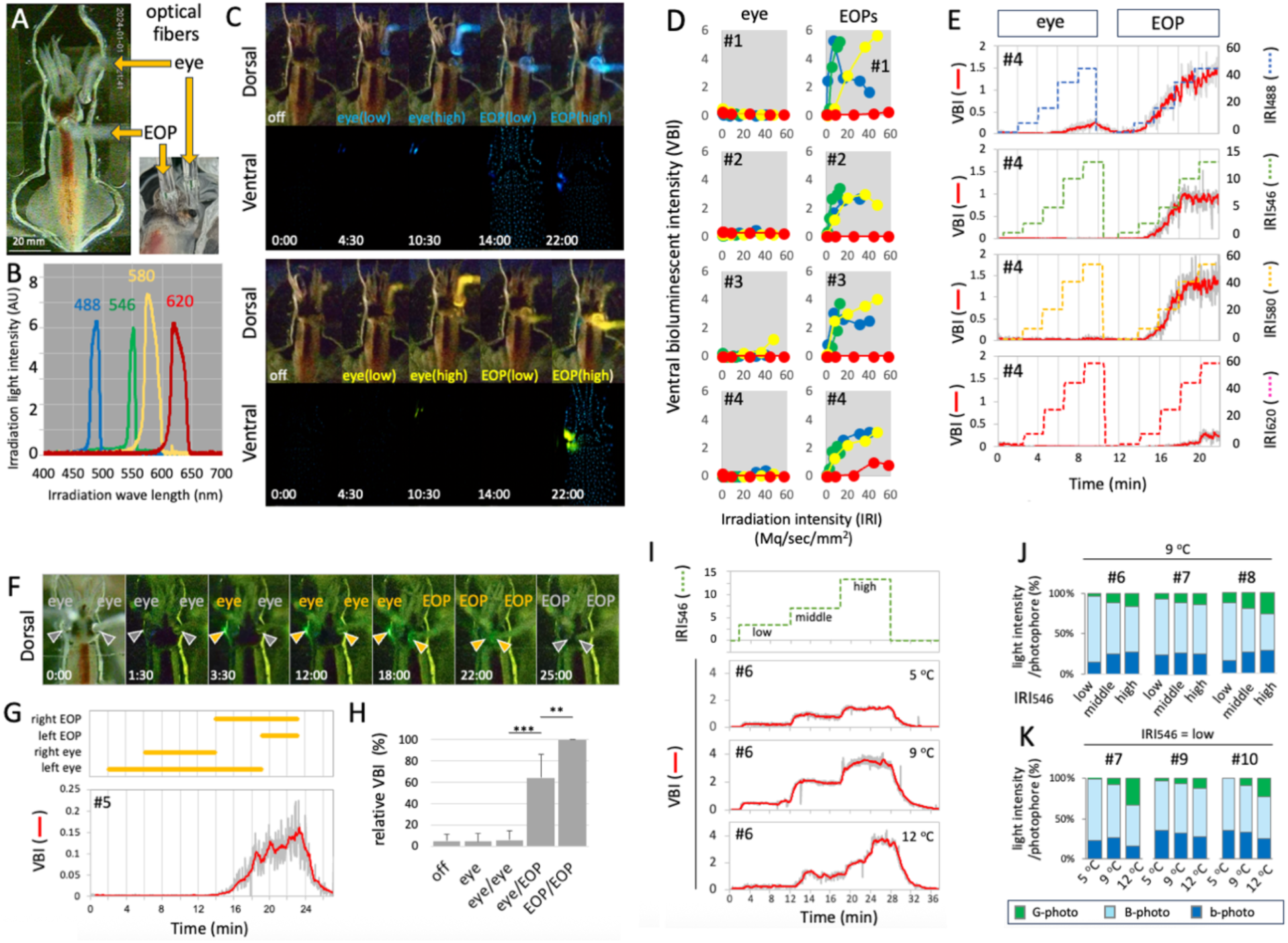
Functional characteristics of ventral illumination in response to various light stimuli, light intensities, irradiation points and temperature. (A) A dorsal view with the optical fibers positioned to the right eye and right EOPs. (B) The spectra of light sources used for directed irradiation. (C) Representative still images of synchronous videography capturing an animal’s dorsal and ventral sides with blue-light (*top* panels) and yellow-light (*bottom* panels) irradiation. (D) Correlation between irradiation intensity and ventral bioluminescence intensity with different light sources. (E) Representative frame-by-frame analysis for ventral illumination dynamics in response to a step-wise increase of irradiation intensity at the eye, followed by EOPs. (F-H) Combinations of direct irradiation at the eye and EOPs, and the resulting bioluminescence output. (I) Representative ventral bioluminescence dynamics by stepwise green-light irradiations at different temperatures. (J-K) Relative contribution changes of bioluminescence from three types of photophores (G-photo, B-photo and b-photo) in response to increasing irradiation intensity (J) and water temperature (K). # indicates individual ID. Mann–Whitney U-test (in H): **, p < 0.01; ***, p < 0.001.

## Discussion

Despite being ‘well-known’ examples of suspected counterillumination, there have been only a handful of experiments on this phenomenon in squids (but see (29) for bacteriogenic bioluminescence). A major obstacle is the difficulty of obtaining bioluminescent mesopelagic squids alive and in good condition. However, due to its unique ecological, geographical, and food-cultural features, *W. scintillans* in Toyama Bay has long provided rare opportunities to investigate the mechanisms of bioluminescence. Nevertheless, there has been no report demonstrating counterillumination in this species (27). The present study used a state-of-the-art, ultra-sensitive CMOS video camera in conjunction with convenient mirror-reflected (Fig. 2A) and direct (Fig. 4A) ventral viewing methods. The results clearly showed that counterillumination, as first demonstrated by Young in Hawaiian squids (14), also occurs in *W. scintillans*, supporting the long-attested assumption that mesopelagic bioluminescent squids in general are probably all capable of performing true counterillumination.

Unlike other bioluminescent squids, *W. scintillans* has subsets of green-emitting ventral photophores (24, 30, 31). Chemical and microscopic studies of the light-sensing optical system have revealed that squids may be able to distinguish green wavelengths from blue, perhaps for communication with conspecifics (28), an assumption that awaits empirical evidence. Mesopelagic organisms undergoing vertical migrations are expected to adjust their emission spectra in response to ambient light conditions in their ecological environment, because light penetration depth in the sea depends on wavelength, with green light penetrating less effectively than blue (1, 32–34). Consistent with this reasoning, color changes in emitted light were observed to occur with changes in temperature (and therefore water depth) in *Abraliopsis sp.* and *Abralia trigonura.* Although the mechanism of this color change has yet to be fully understood, it has been postulated that individual photophores change the color of their emitted light (16) or that recruitment of different photophore types might be involved (17, 18). Similarly, previous research on *W. scintillans* has yet to address color change in any of the photophores.

The present study found no evidence of a significant color shift in the counterilluminating photophores (Fig. 3F). Instead, emission from G-photo was temperature-sensitive (Fig. 5K, Fig. S7), supporting the hypothesis that a shift in overall ventral emitted hue from bluish to greenish would require changes in the relative contributions of different photophore classes (17, 18). However, the alternative hypothesis, which posits a color shift within individual photophores, remains unresolved. This is particularly relevant given that photophores in deep-sea lanternfish can alter light from blue to green through light interference with a multilayer inner reflector (22), and such structures are present in all three types of ventral photophores (G-photo, b-photo, and B-photo) in *W. scintillans* (Figs. 1H, 1I, 1K,Fig. S2C). The physiological mechanisms underlying the temperature-sensitive illumination of G-photo warrant further investigation into their regulation, whether by hormonal or neuronal control (35, 36). Furthermore, these findings provide a foundation for exploring the mechanism of ambient light adaptation, advancing knowledge of both intensity and color matching (16).

Curiously, G-photo remained illuminated for an extended period in the dark, which does not align with the expected function of crypsis. Further analysis using improved imaging systems will be necessary to accurately estimate the emission spectra and spectral changes of nearly 1,000 photophores scattered across the entire body. Furthermore, to validate the crypsis hypothesis, there is still a lack of evidence meeting the criteria for genuine counterillumination: (1) the intensity of bioluminescence should be correlated with or well matched to background brightness; (2) when viewed from below, the silhouette of *W. scintillans* should appear obscure or less conspicuous; (3) high-contrast illumination patterns should have an obscure outline, providing disruptive coloration (4, 37). As a result, the probabilities of being targeted, attacked, and eaten by predators should decrease. In addition, because *W. scintillans* swims in schools, background matching could be achieved in large clouds, making precise illumination control for each individual squid less important than when swimming alone. Also, the foraging and hunting methods of potential predators, as well as their visual acuity, could affect the illumination strategy of light-emitting squids (38).

Light shielding is another feature that many luminous animals possess to effectively utilize bioluminescence (8, 39, 40). In *W. scintillans*, sudden quenching of light emission from the three photophores at the tips of arms IV can be achieved by covering them with the specialized chromatophores with which they are associated (41, 42). These photophores are presumably used to startle or dazzle predators when the arm-tip photophores are suddenly uncovered. In the present study, irregular and dynamic patterns of light emission were observed from the five subocular photophores. These patterns were partly a result of light shielding by the dermal chromatophores, as presumed for other deep-sea squids (19, 43). This also raises the question of whether suddenly shielding the light beneath the eyes might help with counterillumination or serve other roles, such as signaling (41, 44–46).

Finally, our experimental results demonstrate that extraocular photosensitive vesicles (EOPs), rather than eyes, play a primary role in counterillumination (Fig. 5). This finding aligns with, but also partially differs from, Young’s observation that in the Hawaiian species, both EOPs and eyes contribute to the counterilluminating response (15). The observed inconsistency may result from differences in animal condition, experimental design, or evolutionary divergence (see Fig. 2 in (15)). The latter is particularly noteworthy, as the retina of *W. scintillans* possesses three distinct visual pigments, whereas most other squids have only one type (47), supporting the hypothesis that the firefly squid have two-color vision that may be used for intraspecific communication (28). In addition, all other body parts, excluding EOPs, play little or no role in counterillumination, discounting the possible role of photoreception in the skin in this process (10, 48). Future research should clarify the specific ecological functions of the eyes and retina, such as chromatophore-based light shielding (43), mating signal perception (49), collective vertical migration, and mass stranding ("minage" in Japanese), which predominantly occurs at the new moon.

In summary, although this study identified conditions that trigger synchronous illumination, the unexpected observation of asynchronous illumination across all three photophore types (oc-photo, B/b-photo, and G-photo) offers new insights into the physiological, biological, and ecological functions of photophores in deep-ocean squids.

## Materials and Methods

### Animals and video recording

Fresh *W. scintillans* were obtained live from commercial firefly squid fishing in Toyama Bay, Japan, in March-May, 2025-2026. They were then carefully transported in dark containers to Uozu Aquarium, where they were held in a stock tank for 24 h at 2 °C (the temperature at a depth of around 300 m in Toyama Bay, where the squid dwell during daylight). Subsequently, most specimens were illuminated under dimly lit conditions. The animals were transported to a polystyrene reserve tank (400 x 500 x 300 mm) at the desired water temperature (5-15 °C) to acclimate them under experimental conditions for at least 30 min.

For counterillumination experiments, we developed two systems. The first system is suitable for observing counterillumination in squids under free-swimming conditions with diffuse overhead lighting, while the second system enables directed illumination of targeted areas with different light sources and intensities. In the first system, each individual was placed in an acrylic tank (310 x 60 x 155 mm, thickness: 5.0 mm) filled with aerated seawater at 9 °C (approximately the sea surface temperature in March in Toyama Bay). The ventral photophores were observed from the bottom of the tank through a mirror angled at 45°, and images were recorded using an ultra-sensitive full-HD CMOS camera (NC-H100, NEC, Tokyo) with a zoom lens (LAOWA 100 mm F2.8 2X Ultra Macro APO, Venus Optics, Hefei, Anhui, PRC; with a Nikon FW-ENG Converter TMW-B1) and displayed on a monitor. Dim overhead lighting was provided by a 25-lm white LED lamp (CP-195DB, GENTOS, Tokyo) and a reflective sheet positioned over the top of the tank, creating downwelling, scattered dim light of approximately 0.01 lx. Video images of individual photophores were obtained when the animal was hovering gently at the bottom of the tank, with a focus on the ventral head region, which shows minimal movement during the mantle’s respiratory movements.

In the second system, the animal was placed in a bisymmetrical, two-piece ultra-fit acrylic chamber that allowed minimal movements but did not restrict rhythmic fin movements and mantle contractions for breathing. The chamber was set in an acrylic tray (300 x 300 x 50 mm, thickness 5.0 mm) with thermoregulated inflow and outflow of water. The tray rested on a vacuum-insulated glass plate (300 x 400 mm, thickness 6.2 mm) to prevent condensation and ensure clear ventral views. The apparatus was housed in an aluminum frame with two stacked blocks (1060 x 560 x 2000 mm), covered with blackout curtains in a dark room. Two supersensitive video cameras with monitors were mounted: one (NC-R550, NEC, Tokyo, for dorsal view) above the tray and the other (NC-H100, for ventral view) below it. Electroluminescent (EL) sheets (white, blue, green, red) with long-pass or short-pass filters served as light sources, and a voltage converter adjusted intensity. Targeted point illumination was delivered via optic fibers (2.0 mm diameter, 1 m) mounted on manual three-axis manipulators (NARISHIGE, Tokyo), with positioning adjusted by observing a top-view monitor in the dark.

### Estimation of the bioluminescence intensity of individual photophores

A series of still images with minimal motion blur was extracted from video footage (captured with the first system) and analyzed using the ImageJ v2.1.0/1.53c/Fiji plugin. In the second system, all video frames (60 fps) were converted to TIFF images using DaVinci Resolve 21.0.1. The values of each split RGB color component (8-bit) were measured within the area of each photophore (10 photophores per photophore type). The relative luminescence intensity (photon counts) of each photophore type was estimated as follows. Calibration curves were generated between standard light intensity and the corresponding RGB color value obtained as mentioned above. It is already known that blue light (λmax at 470 nm) and green light (λmax at 530 nm) are emitted from *W. scintillans* photophores (25). These wavelengths of light were generated in standard intensity series by a xenon lamp equipped with a Φ100 mm optic fiber cable and a diffraction grating unit. The intensity of light transmitted through the optic fiber was measured directly with a spectrophotometer (USB4000, Ocean Optics, Dunedin, FL, USA) using OPwave+ (Ocean Photonics). Calibration was done with a Tritium light source (Beta Light Source, Saunders-Roe and Nuclear Enterprises, UK) (50, 51). The terminal surface of the fiber was then filmed perpendicularly with a super-sensitive CMOS camera, using the same image acquisition conditions as those captured in the counterillumination experiments. Different light intensities from the xenon lamp were achieved using a series of ND filters (Fujifilm). Changes in luminescence intensity were expressed as photon counts/s/pixel, or as fold change after 1 min of overhead LED lighting turned on (FCoff/on) or off (FCon/off). Statistical analyses were performed in the R package (v.4.4.1) and RStudio.

### Histology and Transmission Electron Microscopy

Animals were anesthetized using 30 mM MgCl_2_ in seawater prior to euthanasia. The dermal skin tissue containing each photophore was isolated and fixed with 4% paraformaldehyde containing 1% glutaraldehyde in phosphate-buffered saline (PBS) for 24 h, then replaced with PBS at room temperature. Paraffin sections were stained with Hematoxylin-Eosin (HE-stain), Elastica van Gieson (EV-stain), Masson Trichrome (MT-stain) and Klüver-Barrera (KB-stain) according to standard procedures. For TEM, samples were pre-fixed with 2.5% glutaraldehyde in 30 mM HEPES buffer (pH 7.4) containing 100 mM NaCl, 2 mM CaCl_2_ and 0.45 M sucrose for 24 h at 4°C, and post-fixed with 1% osmium tetroxide and 10 mM potassium ferricyanide in 30 mM HEPES buffer (pH 7.4) containing 100 mM NaCl, 2 mM CaCl_2_ for 1.5 h at room temperature. The following steps, including dehydration, embedding, ultrathin sectioning, electron staining, and imaging, were carried out as previously described (52, 53).

All experiments were carried out with permission from the Head of Uozu Aquarium, Uozu City, Toyama Prefecture, Japan. This study was approved by the Animal Care and Use Committee of Shimane University (permit no. MA2-2, MA7-06).

## Acknowledgments and funding sources

We thank the members of Uozu Fisheries Cooperative, Mitsuwa Setnet Cooperative, and Uozu Aquarium for providing healthy squids. We are grateful to Shirou Nukiwa (Mitsuwa Setnet Cooperative), Takayuki Sato (NEC Platforms, Ltd.), and Jérôme Mallefet for access to their facilities and equipment, and to Kyohei Kudamatsu, Aoi Shinoda, Shiori Umeda, Miyuu Minemura and Kei Nakano for assistance with experiments and data analysis. Funding was provided by the Faculty of Life and Environmental Science at Shimane University and Kakenhi (22H05681, 25K02088, 25K09247) to N.H. We also thank Ian Gleadall for critically reading the manuscript. This research was conducted in part as the SDGs Research Project at Shimane University.

## Supporting Information

**Figure S1.**
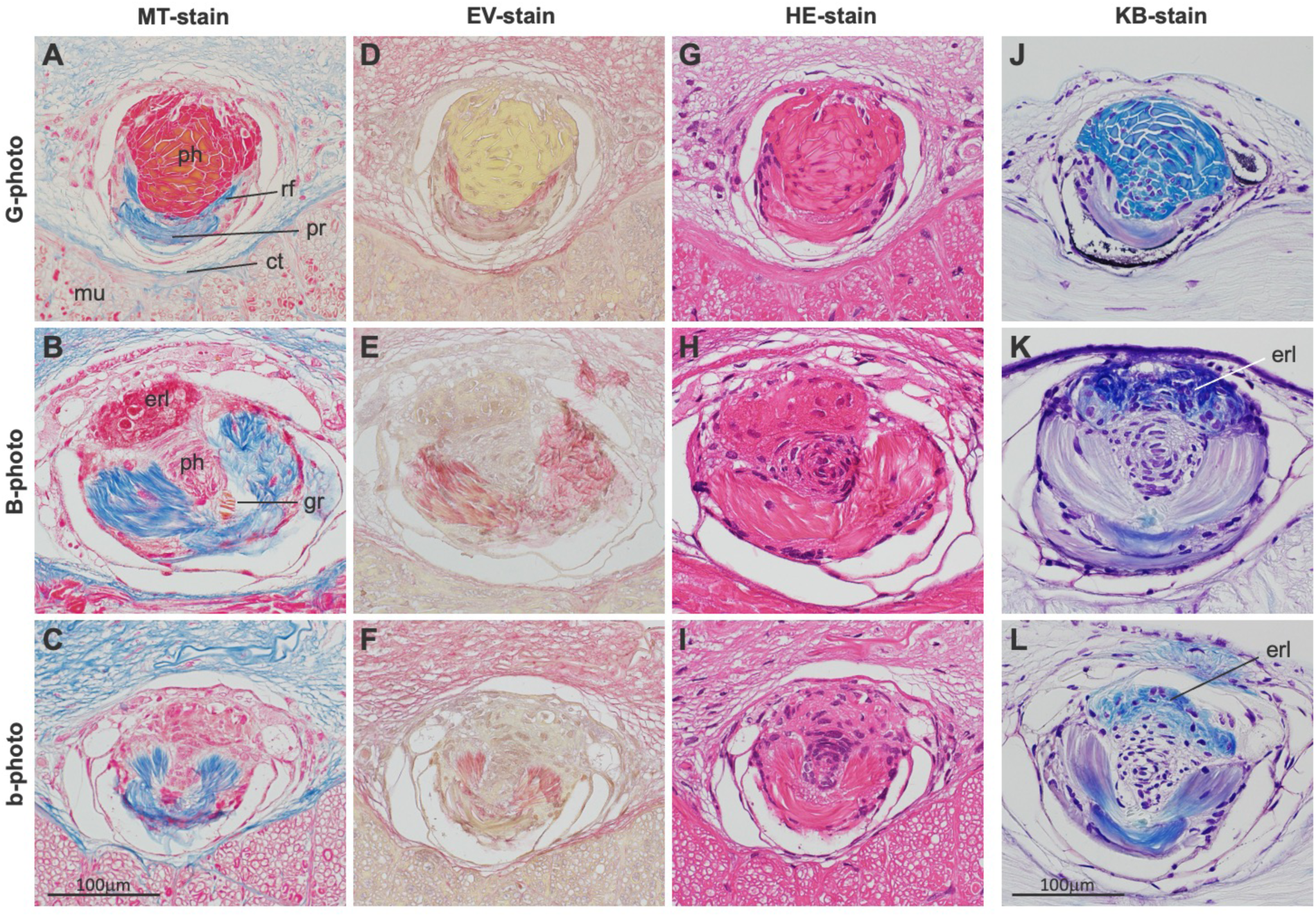
Histology and TEM of the photophores. The longitudinal paraffin sections of each type of photophore (*G-photo, B-photo* and *b-photo*) are stained with Masson Trichrome (*MT-*stain, A-C), Elastica van Gieson (*EV-stain,* D-F), Hematoxylin-Eosin (*HE-stain,* G-I), and Klüver-Barrera (*KB-stain,* J-L). B-photo and b-photo are morphologically similar and indistinguishable in their structural components (also see Figs. S1 and S2). Both photophores contain Orange G-positive granules (*gr*) with high electron density (see Fig. S2C) located between photocytes and the posterior reflector. Only G-photo exhibits the photocytes in yellowish with EV stain (panel D). Both B- and b-photos exhibit positive KB staining in the region corresponding to endoplasmic reticulum lamellae (erl) overlaying the photocytes (panels K and L). ph: photocytes; erl: endoplasmic reticulum lamellae; rf: reflector; pr: posterior reflector; ct: connective tissue; mu: muscle; gr: granules;

**Figure S2.**
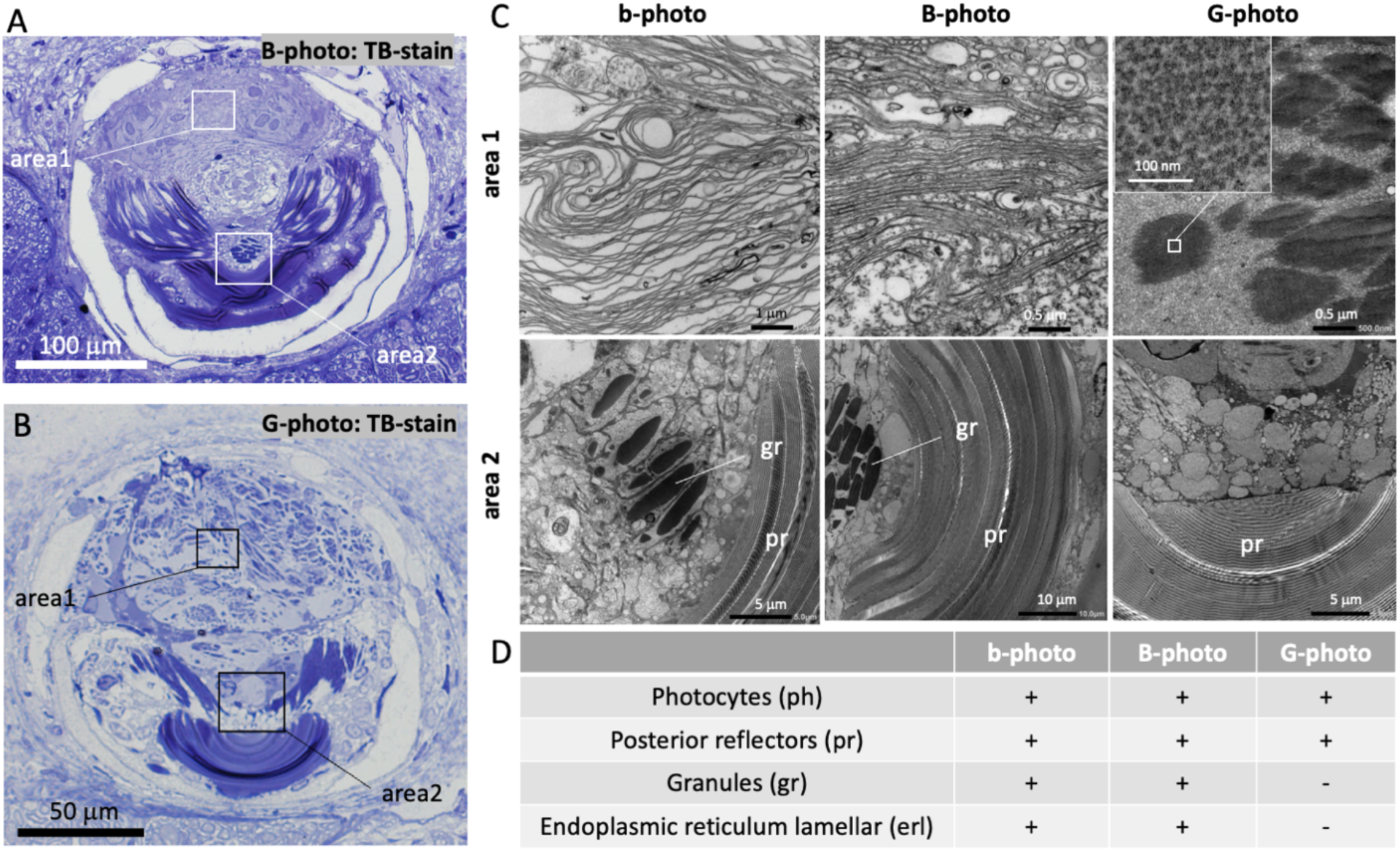
Transmission electron microscopy of the luminous organs in *W. scintillans*. A-B, Representative photographs of B-photo (A) and G-photo (B) stained with toluidine blue (TB-stain). **C,** The photographs present transmission electron micrographs of the boxed regions highlighted in panels A and B. It is noteworthy that while B- and b-photos exhibit a lamellar pattern of membranous structures in area 1, this structure is absent in G-photo. Instead, G-photo displays a dense array of well-aligned triangular rods (C, *inset*, *top-right* panel). **D,** Summary of the presence or absence of structural components in each photophore type, as validated by multiple staining methods.

**Figure S3.**
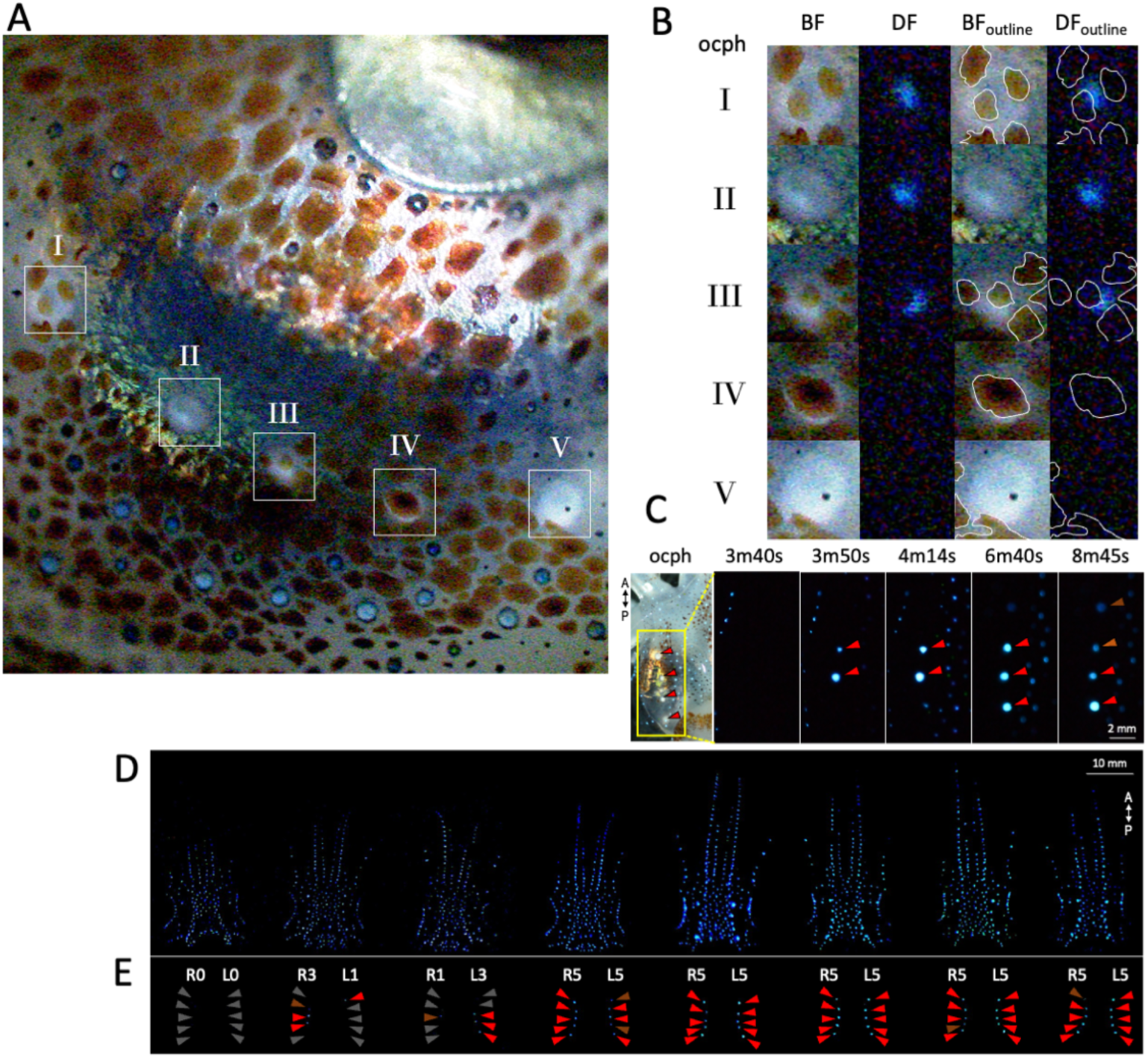
Illumination of ocular photophores. (A) A close-up view of the ventral left eye rim having five ocular photophores (oc-photo) in a line (*I-V*). (B) Skin chromatophores overlay oc-photo and can partially or completely shield their light emission. There was no light emission from oc-photo V despite the absence of a chromatophore overlay. BF, bright field; DF, dark field; retouched outlines clearly demarcate the chromatophores. (C) Representative time-lapse images of light-emitting oc-photo (*red arrowheads*). A, anterior direction; P, posterior. (D, E) Free-swimming individuals (n=8) showing different illumination patterns in oc-photo. Ventral views of the anterior region of arms and head, and the corresponding oc-photo viewed in isolation. *Arrowheads* point to each oc-photo coded according to emission state: off (*gray*), weak (*dark-red*) or bright (*red*). Rx, right; Lx, left. Numerals (x) with R and L indicate the number of emitting photophores on each side.

**Figure S4.**
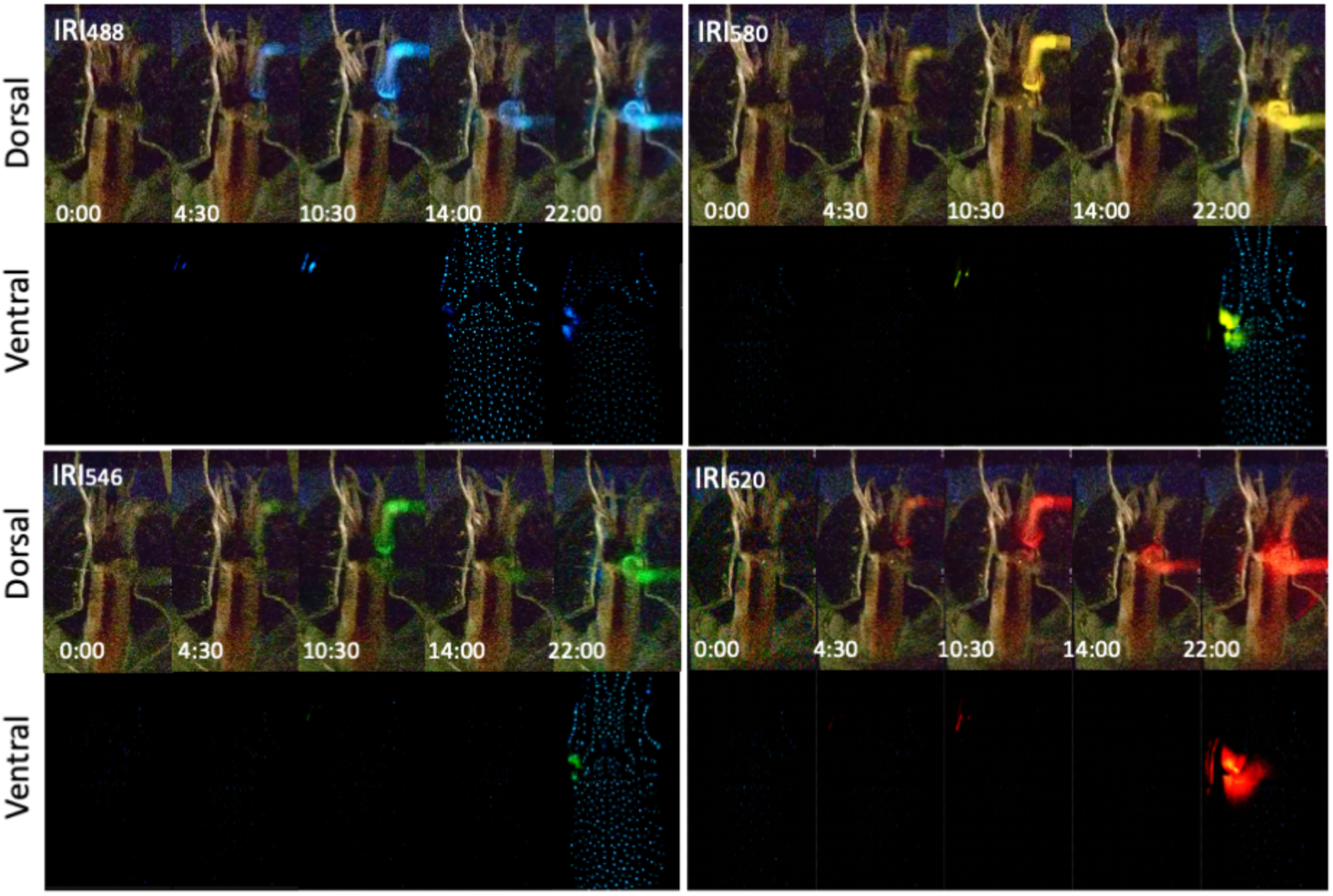
Synchronized video-recording with two supersensitive video cameras in counterillumination. Time-course of counterillumination responses upon directed irradiation of blue (λ_max_ 488 nm), green (λ_max_ 546 nm), yellow (λ_max_ 580 nm) and red (λ_max_ 620 nm) light to either right eye or right EOPs. Note that strong irradiation causes light to penetrate the animal’s body or leak from its periphery. Time code indicates min:sec.

**Figure S5.**
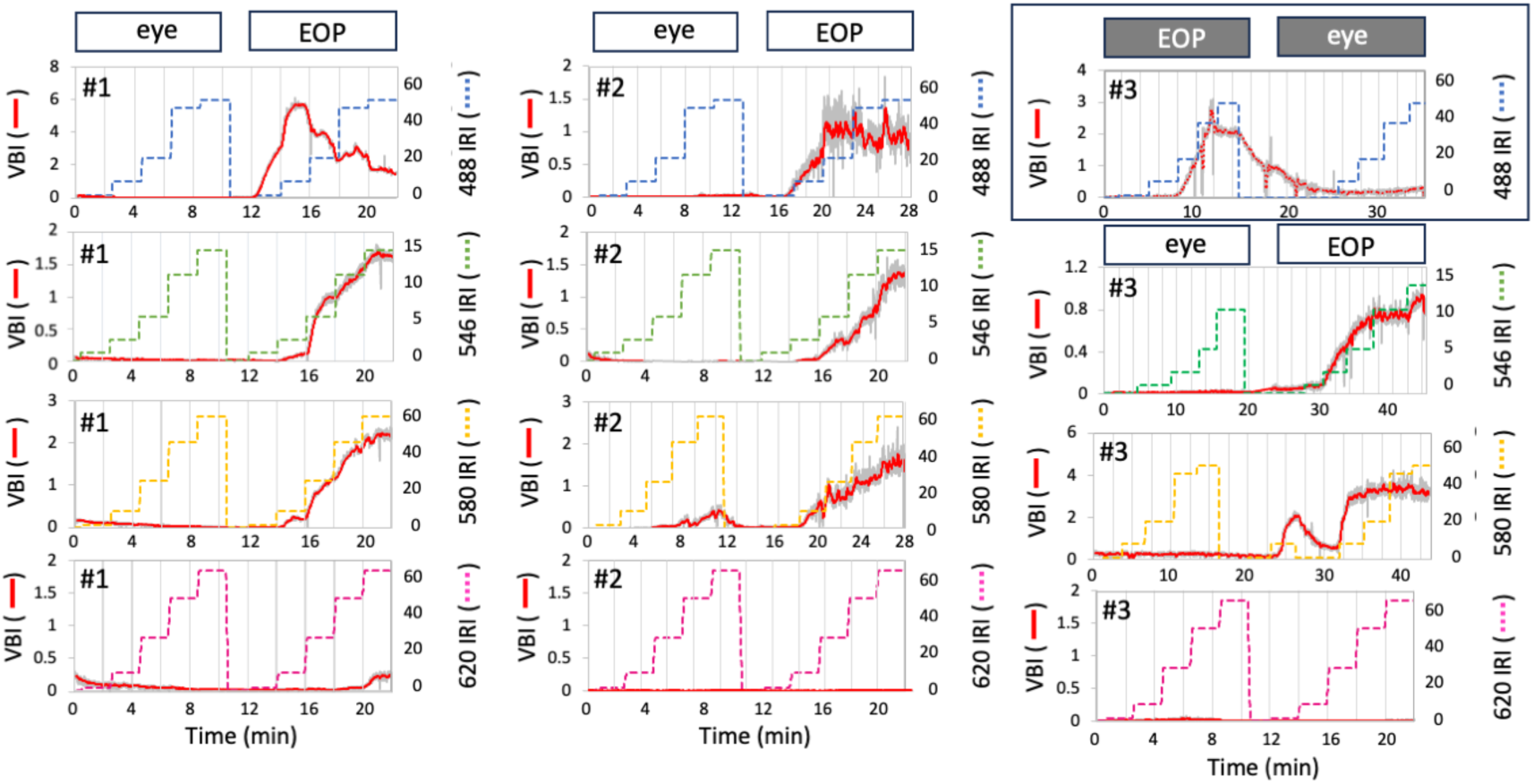
Counterilluminating responses elicited by applying directed irradiation to the eye or extraocular photoreceptors (EOPs) using various wavelengths and intensities. The average illumination intensity from the lower half of the ventral body was quantified by measuring mean gray values after converting videos to the TIFF images at frame-by-frame resolution. Light intensity was calibrated with a standard tritium light source coupled with a series of ND filters (*Materials and Methods*). Simultaneous, real-time monitoring of dorsal irradiation allows us to validate the precise positioning of the irradiation point and its adjustment. Shown are the data from three animals (#1, #2, and #3) and four different light sources.

**Figure S6.**
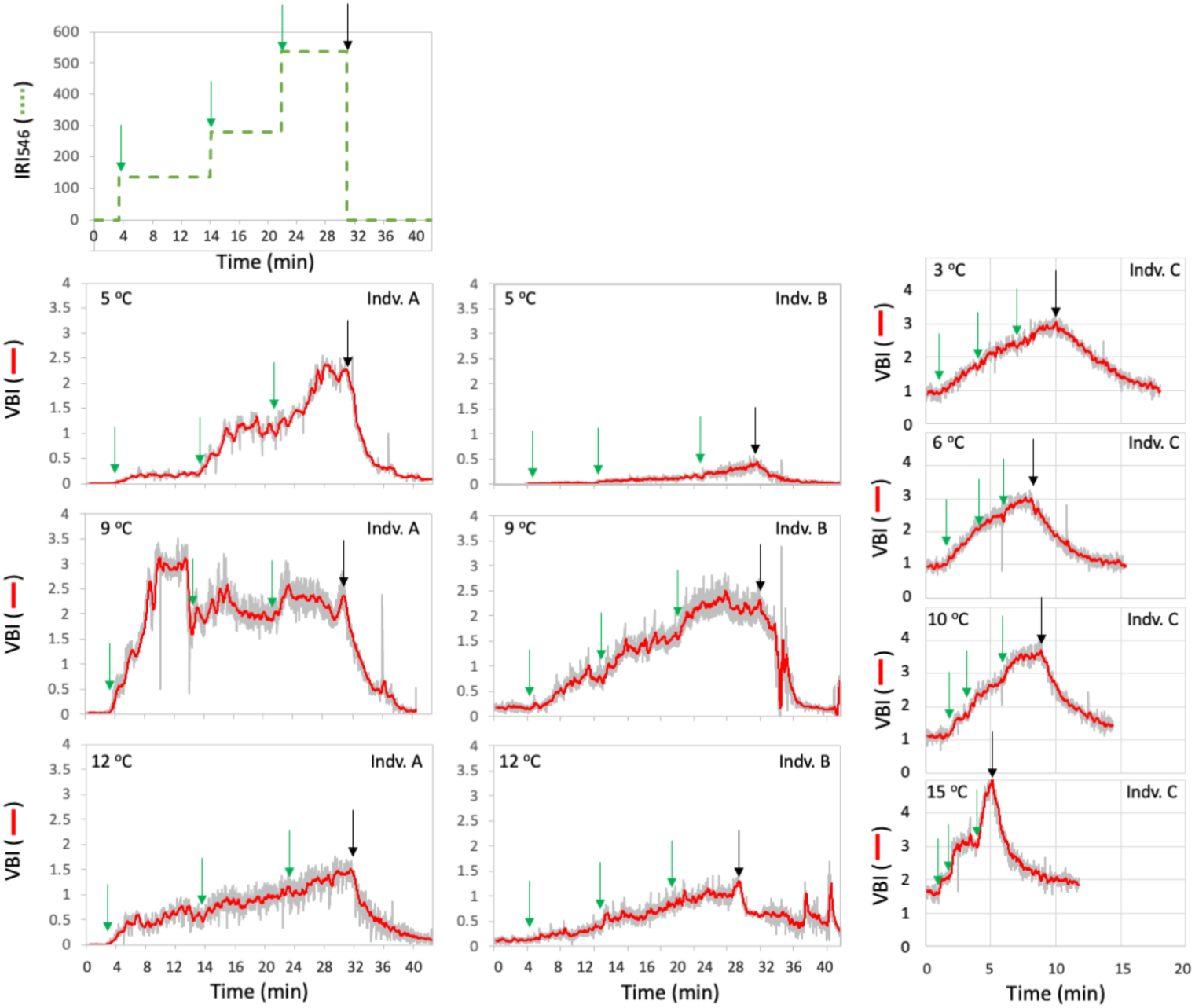
Ventral bioluminescence dynamics by stepwise green-light irradiations at different temperatures. A green light at λ_max_ 546 nm was applied to EOPs on the right side of the animal with a stepwise increase in intensity (9.5, 19.8, 37.8 Gq/sec/mm^2^). Each frame from the video clips (60 fps) was converted to a TIFF image, and the mean gray value of the region of interest (ROI) corresponding to the lower mantle was measured. This region was chosen because it exhibited minimal movement and lacked light contamination from the optical fiber.

**Figure S7.**
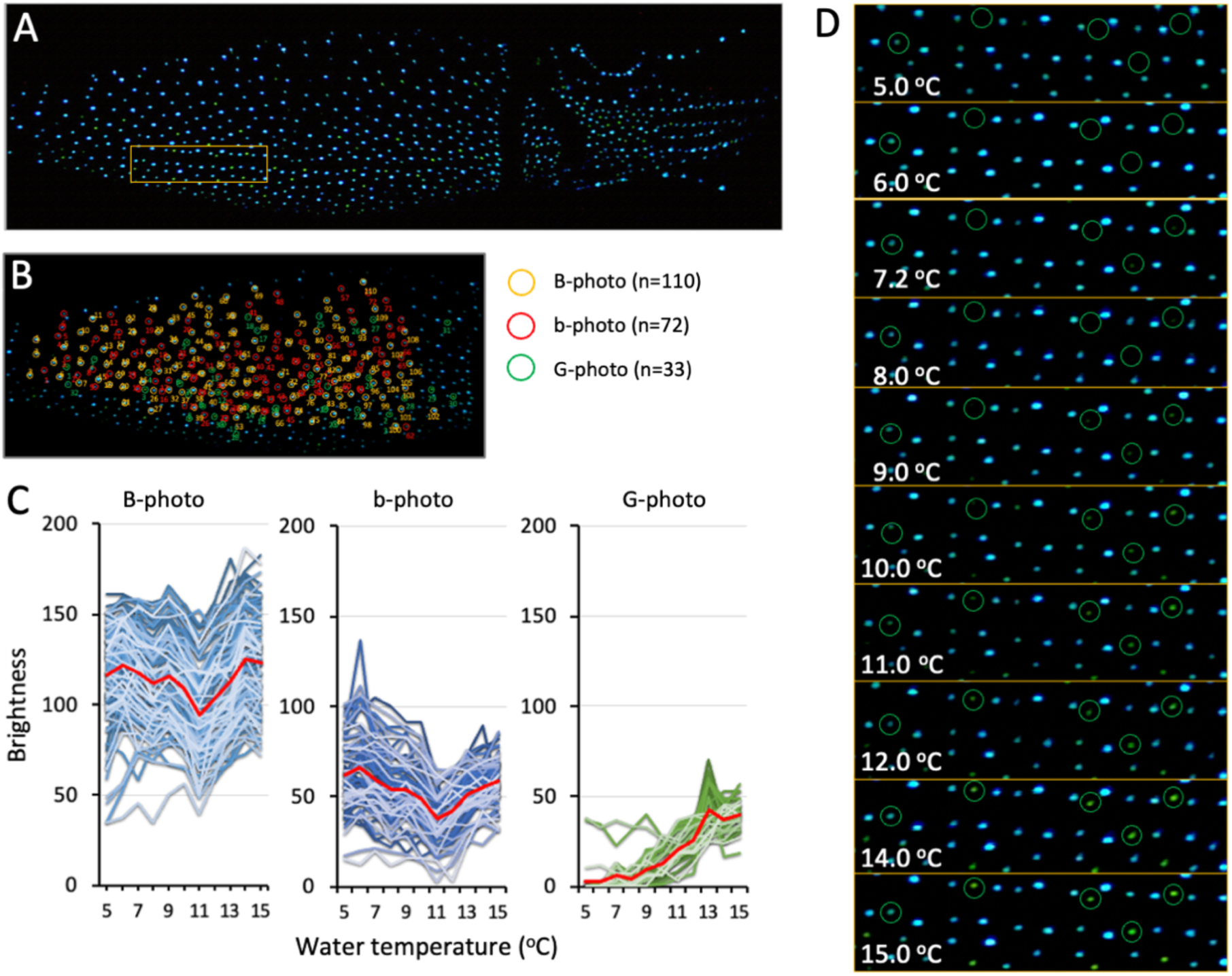
Temperature dependence of illumination in G-photo. A) A whole animal view from the ventral side illuminating most of the photophores at 15.0 °C in darkness. (B) The same image for every single photophore, with each given an ID number and marked in a color corresponding to its type (B-photo, b-photo, or G-photo). The photophores that were ambiguous for typing or less visible due to peripheral localization were eliminated from the analysis. (C) Changes in brightness (as estimated by gray value measurements) of every single photophore (and the average shown in red) were plotted as a function of water temperature. (D) Photographs of the selected area (a *broken box* in (A)) showing G-photo (*green* circles) that were least illuminated at 5.0 °C and became illuminated as the temperature increased.

### Movie S1 (separate file)

This video shows two cycles of counterillumination in the ventral and ocular photophores, with bioluminescence increasing and then decreasing when exposed to dim light from the dorsal side. The video is viewed from the ventral head region and is played at 60x speed.

### Movie S2 (separate file)

This video shows a gradual, steady increase in ventral luminescence in response to four brief, intermittent exposures to dim light, followed by the extinguishing of bioluminescence in continuous darkness. The video is played at 10x speed.

### Movie S3 (separate file)

This video shows only an increase in bioluminescence after prolonged exposure to dim light. The animal quickly rushed out of view. The video is played at 5x speed.

### Movie S4 (separate file)

This movie shows a dorsal view of the squid placed in a semi-closed chamber, with green light directed at both the eye and the extraocular photosensitive vesicles.

### Movie S5 (separate file)

This movie shows a ventral view of the squid and demonstrates how its photophores adjust illumination intensity based on the location and strength of directed light on specific dorsal body regions. The video is played at 120x speed.

### Movie S6 (separate file)

This movie shows a zoom-in view of a part of the ventral surface (Fig. S7D) where the green photophores become illuminated, while the blue photophores remain lit as water temperature increases. The video is played at 1000x speed (duration: 80 min.).

## Contact and competing interest information for all authors

Contact to Noritaka Hirohashi

The authors declare no competing interests.

